# Synergistic effect of short- and long-read sequencing on functional meta-omics

**DOI:** 10.1101/2021.04.22.440869

**Authors:** Valentina Galata, Susheel Bhanu Busi, Benoît Josef Kunath, Laura de Nies, Magdalena Calusinska, Rashi Halder, Patrick May, Paul Wilmes, Cédric Christian Laczny

## Abstract

Real-world evaluations of metagenomic reconstructions are challenged by distinguishing reconstruction artefacts from genes and proteins present *in situ*. Here, we evaluate short-read-only, long-read-only, and hybrid assembly approaches on four different metagenomic samples of varying complexity and demonstrate how they affect gene and protein inference which is particularly relevant for downstream functional analyses. For a human gut microbiome sample, we use complementary metatranscriptomic, and metaproteomic data to evaluate the metagenomic data-based protein predictions. Our findings pave the way for critical assessments of metagenomic reconstructions and we propose a reference-independent solution based on the synergistic effects of multi-omic data integration for the *in situ* study of microbiomes using long-read sequencing data.

## Background

Third-generation, single-molecule, long-read (LR) sequencing is considered to be the next frontier of genomics [1], especially in the context of studying microbial populations [2,3]. Given the ability to attain read lengths in excess of 10 Kbp [4] and sequence accuracy improvements [5], LR sequencing has been recommended for its ability to resolve GC-rich regions, complex and repetitive loci, and segmental duplications in genomes [4]. However, LR applications to study microbiomes have focused on genome assemblies [6,7], closing a select few bacterial genomes [8], haplotype and strain resolution [9] as well as mock (low diversity) communities [3]. Stewart *et al.*, recently were among the first to demonstrate the utility of using LRs for improving upon existing protein databases owing to a large collection of novel proteins and enzymes identified [10], thereby hinting at the benefits of LR also for functional microbiome studies.

Single base-accuracy of raw LRs remains lower - for now - compared to short-read (SR) methodologies [11]. Several approaches including assembly-based and/or including polishing steps have been developed [11–13] to increase the accuracy. The impact of remnant errors in LR assemblies on gene calling and thereby protein prediction was recently highlighted by Watson *et al*. [14]. Hybrid (HY) assembly methods [15,16] using both SRs and LRs have been proposed to further reduce the error rates compared to LR-only assemblies. While Watson *et al*. [14] showed that insertions/deletions (indels) play a critical role in microbial protein identification, the overall impact of assembly methods on understanding the functional potential of microbial communities is lacking.

Here, we demonstrate that metagenomic assembly methods (SR, LR and HY) not only differ markedly in their overall assembly performance, but also in the inferred functional potential. We reveal the effects of the assembly method on predicted genes and proteins in samples with a low to high diversity, from mock communities to human fecal and rumen metagenomes. We found proteins which are exclusive to respective assemblers and additionally demonstrate using metatranscriptomic and metaproteomic data available for the human fecal sample the synergistic effect on protein verification. Our results indicate that irrespective of sample diversity, the sequencing and assembly strategies impact downstream analyses and that complementary omics are a key dimension for functional analyses of microbiomes.

## Results and Discussion

To understand how sample diversity, assembly quality, and assembly approach are linked, we assembled published metagenomic (metaG) data from a mock community (Zymo), a natural whey starter culture (NWC), a cow rumen sample (Rumen), and a novel metagenomic dataset from a human fecal sample (GDB) which was complemented with metatranscriptomic (metaT) and metaproteomic (metaP) data. The samples’ diversity ranged from low (Zymo and NWC) to high (GDB and Rumen). As expected [10], the assembly approach affected strongly the quality of the resulting assembly (Supp. Fig. 1). LR and HY approaches generated fewer contigs with a larger N50 value. However, other assembly metrics, e.g., the total assembly length, varied between the samples and assembly types. The metaG read mapping rate (including multi-mapped reads), as a proxy of data usage, was unaffected by the assembler choice when considering all contigs, though the values for the LR assemblies were a bit lower than for SR or HY assemblies of the high-diversity samples (GDB and Rumen). However, the mapping rates dropped markedly in SR assemblies, especially in NWC and Rumen, when filtering out contigs below 5000bps (Supp. Fig. 2). In GDB, we observed higher metaT read mapping rates in SR and HY assemblies than in LR assemblies. This indicates the complementarity of SR and LR data. The mapping rates decreased considerably in SR assemblies when removing short contigs (Supp. Fig. 3) suggesting the presence of expressed genes located on these contigs. This demonstrates the loss of information when contigs below a certain threshold are removed, which is frequently done in metagenomic studies.

Comparing assemblies pairwise, we observed higher dissimilarities between the LR and SR/HY assemblies than within the latter groups. Additionally, OPERA-MS-based HY assemblies clustered together with the SR assemblies on which they were based (Supp. Fig. 4). To assess functional potential overlap between the different assembly approaches, we studied the proteins found in the individual metagenomes. The overall number and quality of predicted proteins was highly influenced by the assembly approach. In highly diverse metagenomes (GDB and Rumen), the total number of proteins in SR and HY assemblies was higher (by a factor of up to 3.67) than in LR assemblies (Fig 1i). However, throughout all samples, the SR and HY approaches produced more partial proteins (incomplete CDS). We clustered the predicted protein sequences and found a considerable number of proteins exclusive to individual assembly. We also found proteins that were shared within a subset of the assemblies only. Furthermore, we observed that increased sample diversity resulted in an overall increase in the number of exclusive proteins (Fig 1ii).

**Fig. 1:**
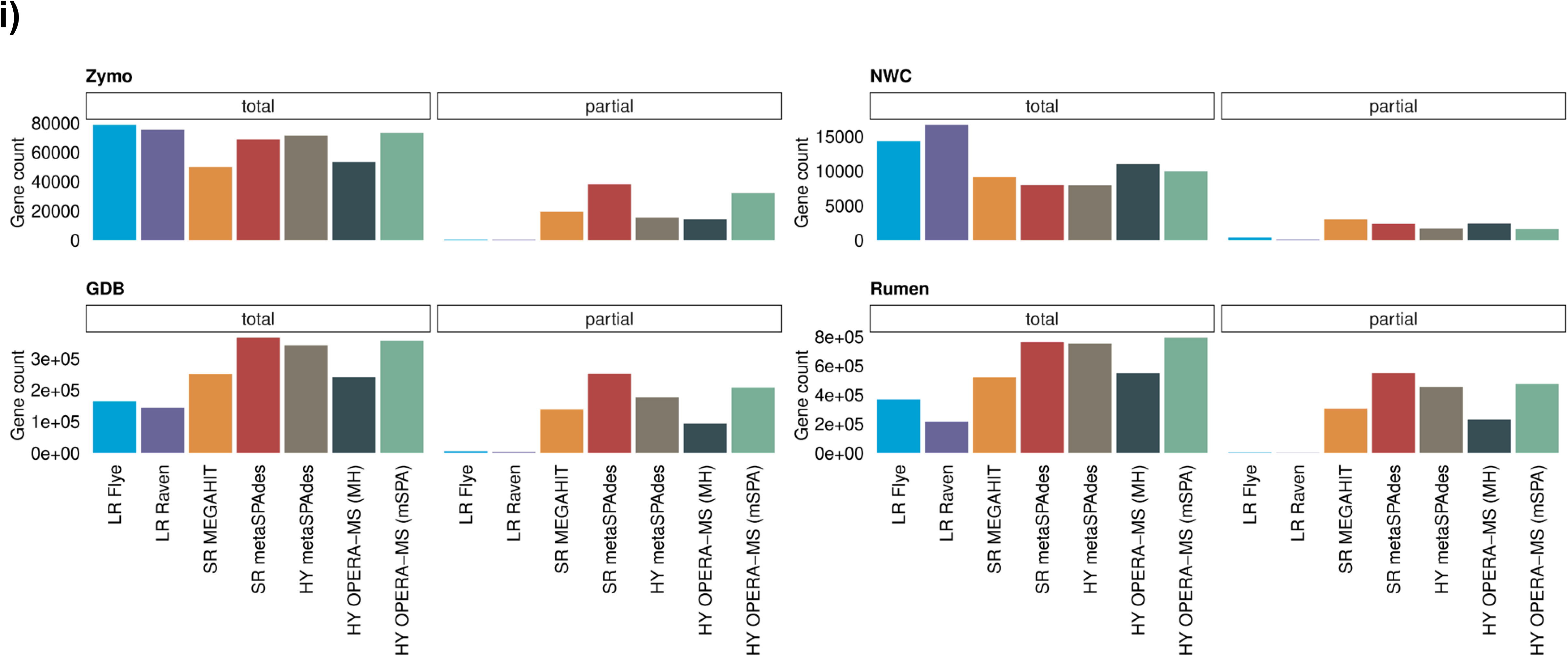
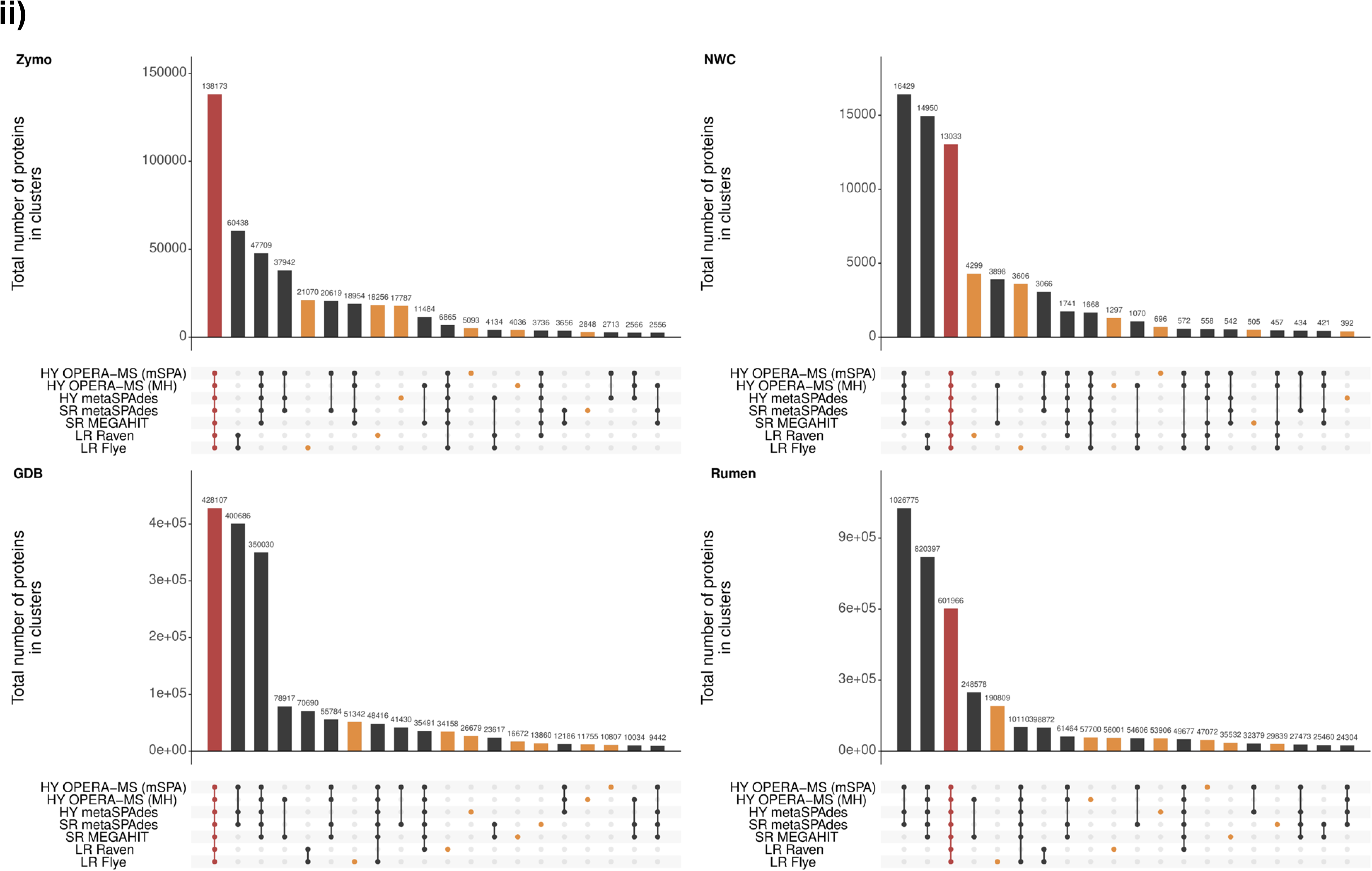
Discrepancy and uniqueness of predicted proteins in assemblies. **i** Number of proteins (total and partial) predicted by Prodigal in each assembly and sample. The color corresponds to the tool used for metagenomic assembly. **ii** Number of shared assembly proteins which were clustered using MMSesq2 per sample. Each protein cluster was labeled by the combination of assembly tools represented by the clustered proteins (i.e., the assembly where these proteins originated from). The depicted number of shared proteins per assembly tool combination is the total protein count over all associated clusters. Top 20 combinations are shown. The number of proteins found in clusters representing all assembly tools is highlighted in red; the number of proteins exclusive to an assembly is highlighted in orange.

As reported previously by Watson *et al.* [14], errors in LR assemblies can have an impact on the predicted proteins. To evaluate how the sample diversity might affect this, we mapped the predicted proteins against the UniProtKB/TrEMBL non-redundant (nr) protein database and computed the query/subject length ratio. In all cases, the density distribution of the ratio values had two peaks (below 0.5 and around 1) though the differences between the assembly methods varied across the samples (Supp. Fig. 5). Considering the above findings and despite multiple rounds of polishing, we cannot disregard the impact of errors in long reads affecting the results. Furthermore, we are aware that the results may also be affected by the sequencing depth and gene prediction methods. One also has to account for the microbial composition per sample, given that a large proportion of proteins from the Rumen sample might not have homologs within the used database.

Due to the differences in annotations, which we found to be exclusive to individual assembly approaches, we subsequently studied the effect of assembler choice on two well defined, functionally relevant classes of genes: ribosomal RNA (rRNA) and antimicrobial resistance (AMR) genes. Overall, the total number of rRNA genes recovered by LR and HY approaches was higher across all samples. Within the archaeal and bacterial domains, LR and HY assemblies led to the prediction of more complete genes compared to SR (Supp. Fig. 6). When analysing AMR proteins and focusing only on “strict” hits (i.e. excluding loose hits flagged as “nudged” by the RGI tool, see Methods), HY assemblers were more adept at identifying these proteins compared to either SR or LR. Moreover, LR assemblies contained more “nudged” hits than SR or HY assemblies, suggesting that error rates or other factors might have affected the reconstruction of some AMR genes (Fig 2i). Interestingly, we did not identify any AMR hits in the NWC metagenome, possibly due to it being a food-grade additive [17]. When comparing the overlap of the Antibiotic Resistance Ontology (ARO) terms covered by “strict” hits, we found that some AROs were only identified in SR and HY assemblies, but not in LR, whereas no AROs were found in LR assemblies only (Fig 2ii).

**Fig. 2:**
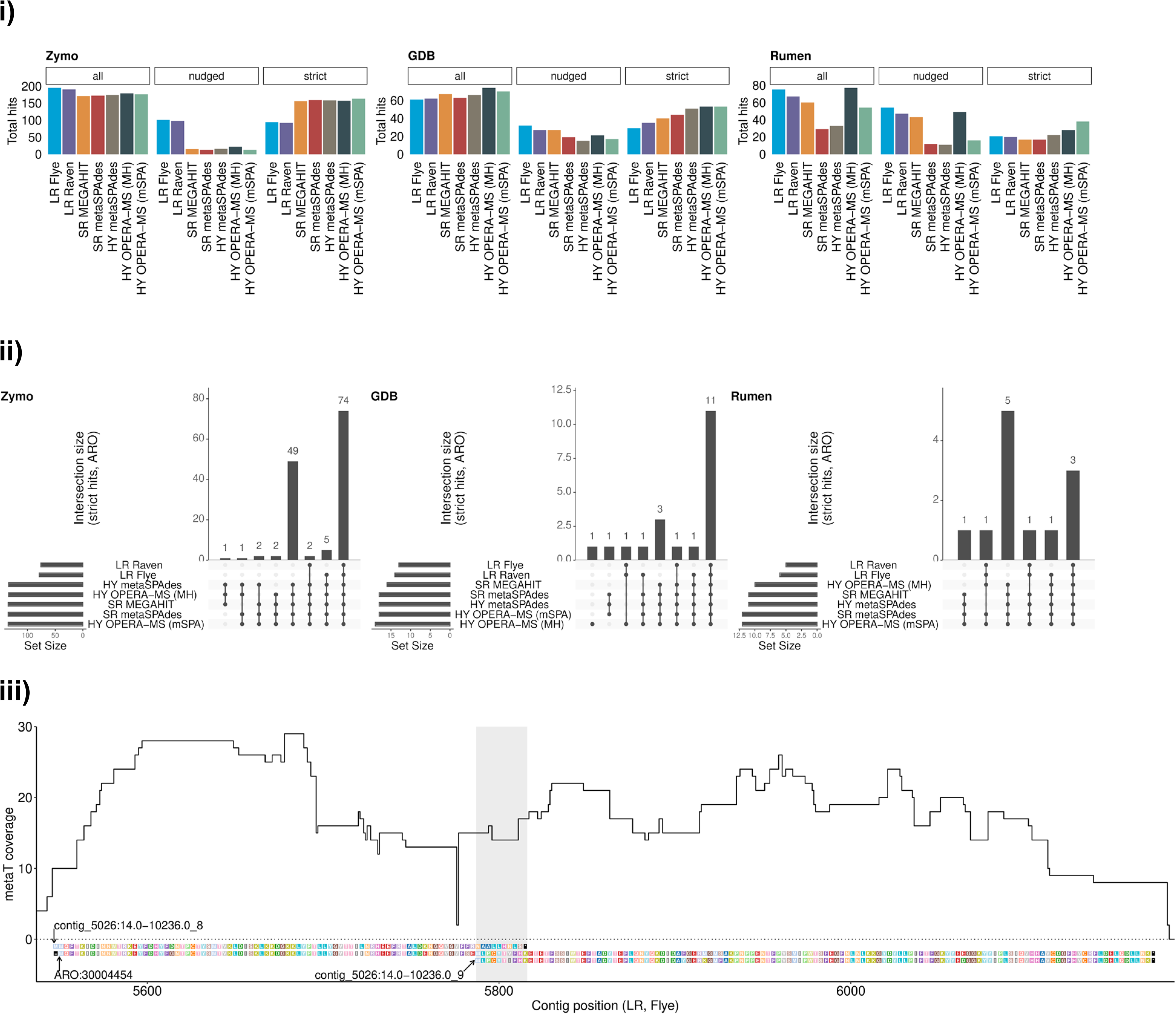
Assembly affects antimicrobial resistance gene identification. **i** Number of hits (total, “strict” and “nudged”) for each assembly and sample when searching the assembly proteins in the CARD database using RGI. Sample NWC was excluded because no hits were found in any of its assemblies. “Nudged” hits are loose hits flagged as “nudged” by RGI; the remaining hits are “strict” hits. **ii** Number of AROs which were covered by “strict” RGI hits by different assemblies per sample. The bar plot shows the number of shared AROs per assembly tools combination. **iii** Metatranscriptomic (metaT) coverage of the two coding sequences (CDSs) from the long-read (LR) assembly constructed with Flye and having a “nudged” RGI hit to ARO 3004454 (a chloramphenicol acetyltransferase) in sample GDB. The x-axis represents the contig coordinates and the y-axis the metaT coverage. The amino acid sequence of the two CDSs and the ARO is included in the plot.

To validate the exclusive AROs found in SR and HY assemblies, we assessed metaT and metaP coverage of the corresponding genes and proteins in the GDB sample. The genes mapping to the exclusive AROs had an average metaT coverage above 14x in the SR and HY assemblies suggesting that these genes are expressed *in situ*; the few “nudged” hits were below 6x (Supp. Tab. 1). However, we did not identify these genes in the metaP data potentially due to low expression levels, variation in extraction protocols, and/or post-translational modifications affecting the peptide/proteomic recovery. Though no “strict” hits were found in LR assemblies, some of their “nudged” hits had an average metaT coverage above 10x. To understand why these seemingly expressed genes obtained only a partial hit, we focused on two “nudged” hits assigned to ARO 3004454 (a chloramphenicol acetyltransferase) in the LR assembly constructed with Flye. We found that the corresponding coding sequences (CDSs) were located on the same contig and had an overlap of 29 bps. The sequence alignments showed that the respective genes represent two fragments of the true CDS (corresponding to ARO 3004454) most likely created by an indel which introduced a frameshift and also a premature stop codon. This finding was also supported by the metaT coverage extending beyond the stop codon of the first CDS until the end of the second CDS with a single drop in coverage before the putative indel (Fig 2iii).

To identify high-confidence proteins without the need for a reference, we first considered proteins and protein clusters found in all assemblies which represented 22.97% of the proteins and 8.54% of the protein clusters. These included genes reconstructed by the different and independent assembly approaches, thus lending mutual support. We then used the complementary metaT data and included all additional proteins with an average metaT coverage >= 10x and the corresponding protein clusters. This doubled the number of high-confidence protein clusters (17.63%) and increased the percentage of high-confidence proteins to 30.32%.

## Conclusions

We reveal that sample diversity, along with assembly-mediated effects influence prediction of genes and proteins. This causes discrepancies between the assemblies, thereby requiring complementary means to validate these predictions. The observed discrepancies included conserved and also functionally relevant genes (rRNA and antimicrobial resistance genes, respectively), potentially impacting phylogenetic as well as functional studies. To overcome this, we propose a reference-independent approach to identify high-confidence genomic reconstructions by combining metagenomic and metatranscriptomic data. Overall, we show that the sequencing approach and assembly strategy can have a significant impact on the characterization of the microbiome’s functional potential and demonstrate the added value of multi-omic strategies for reconstruction quality evaluation, i.e. going beyond their original purpose, to resolve the functional microbiome.

## Methods

Freshly collected human fecal samples from a healthy volunteer (GDB) were immediately flash-frozen in liquid nitrogen and stored at −80 °C; high-molecular weight (HMW) DNA was obtained following the protocol proposed recently [8], with minor modifications; samples were sequenced on Illumina and Oxford Nanopore MinION respectively. Metagenomic sequencing data of three publicly available samples was included: the Zymo mock community (Zymo) [3], a natural whey starter culture (NWC) [17] and a cow rumen (Rumen) [10] dataset. Assemblies were built from short reads (SR), long reads (LR), and short and long reads (HY). The LR and HY assemblies were polished. All assemblies were annotated by predicting rRNA genes and proteins, and matching the latter to the CARD database [18]. For each sample, assemblies were compared, and proteins were clustered. For the GDB sample, metatranscriptomic (metaT) and metaproteomic (metaP) data were additionally used in the downstream analysis. Detailed information on extraction, sequencing and analysis can be found in the Supplementary Information.

## Abbreviations

SR: short reads
LR: long reads
HY: hybrid (approach/assembly)
metaG: metagenomic (data)
metaT: metatranscriptomic (data)
metaP: metaproteomic (data)
AMR: antimicrobial resistance
rRNA: ribosomal RNA

## Declarations

### Ethics approval and consent to participate

This study conformed to the Declaration of Helsinki and was approved by the ethics committee of the Physician’s Board Hessen, Germany (FF38/2016).

## Consent for publication

All authors acknowledge the content of this manuscript and consent to its publication.

## Availability of data and materials

Processed sequencing data of the GDB sample is available under BioProject accession PRJNA723028 (Biosamples: metag_sr: SAMN18797629, metat_sr: SAMN18797630, and metag_lr: SAMN18797631). Metaproteomics data of the GDB sample is available at ProteomeXchange under accession PXD025505. The code used for the analysis is available at https://doi.org/10.6084/m9.figshare.14447553 and supplementary data of relevant results is available at https://doi.org/10.6084/m9.figshare.14447559.

## Authors’ contributions

SBB, VG, and CCL designed the study. SBB and RH performed the biomolecular extractions, while RH performed the metagenomic and metatranscriptomic sequencing. VG, SBB, LdN and CCL analysed the data. BJK performed the metaproteomic analyses. PM, MC and PW provided critical feedback and insights. All authors contributed to the writing and revision of the manuscript.

## Acknowledgements

We are thankful for the assistance of Audrey Frachet Bour, Lea Grandmougin, Janine Habier and Laura Lebrun (LCSB) for laboratory support. The experiments presented in this paper were carried out using the HPC facilities of the University of Luxembourg (https://hpc.uni.lu [19]).

## Supplementary Information: Methods

### Sample origin & collection

Human fecal samples were freshly collected from a healthy volunteer (GDB) and immediately flash-frozen in liquid nitrogen. The samples were stored at −80 °C until they were processed for biomolecular extraction.

### Biomolecular extraction

To obtain high-molecular weight (HMW) DNA, we followed the protocol proposed recently [8], with minor modifications. Frozen stool sample was weighed out in triplicates, to 0.7g and aliquoted into phase-lock gel tubes (Fisher Scientific, Waltham, MA), along with a 4mm stainless steel grinding ball (RETSCH 22.455.0003). The sample was subsequently suspended in 500μL PBS (Fisher Scientific, Waltham, MA) with brief gentle vortexing at 10 second intervals repeated 5 times. Thereafter, 5uL of lytic enzyme solution (Qiagen, Hilden, Germany) was added and the samples were mixed by gentle inversion six times, then incubated for one hour at 37°C. 12μL 20% (w/v) SDS (Fisher Scientific, Waltham, MA) was added followed by 500μL phenol:chloroform:isoamyl alcohol at pH 8 (Fisher Scientific, Waltham, MA). The samples were gently vortexed for five seconds, then centrifuged at 10,000g for five minutes. The aqueous phase was decanted into a new 2mL tube. Next, the DNA was precipitated with 90μL 3M sodium acetate (Fisher Scientific) and 500uL isopropanol (Fisher Scientific). After slowly inverting three times, samples were incubated at room temperature for 10 minutes, followed by centrifugation for 10 minutes at 10,000g. The supernatant was removed, and the pellet was washed twice with freshly prepared 80% (v/v) ethanol (Fisher Scientific). Washing was done by adding 1 ml of 80% EtOH, followed by centrifugation for 10 minutes at 10,000g. The pellet was then air dried with heating for ten minutes at 37°C or until the pellet was matte in appearance, and then resuspended in 100μL nuclease-free water (Ambion, ThermoFisher Scientific, Waltham, MA). To the pellet, 1mL Qiagen buffer G2, 4μL Qiagen RNase A at 100mg/mL, and 25μL Qiagen Proteinase K were added. The samples were then gently inverted three times and incubated for 90 minutes at 56°C. After the first 30 minutes, pellets were dislodged by a single gentle inversion. During the 90-minutes incubation, one Qiagen Genomic-tip 20/G column per triplicate sample was equilibrated with 1mL Qiagen buffer QBT and allowed to empty by gravity flow. Samples were gently inverted twice, applied to columns and allowed to flow through. Three stool extractions (triplicates for each sample) were combined per column. Columns were then washed with 3mL Qiagen buffer QC, where 1 ml of QC buffer was added each time and allowed to drain the column. Next, the column was placed in a new, sterile 1.5 mL Eppendorf tube and the DNA was then eluted with 1mL of Qiagen buffer QF prewarmed to 56°C. The eluted DNA was then precipitated by addition of 700μL isopropanol and incubated at room temperature for 10 minutes, followed by inversion and centrifugation for 15 minutes at 10,000g. The supernatant was carefully removed by pipette, and pellets were washed with 1mL 80% (v/v) ethanol. (Washing = add 1 ml EtOH, centrifuge for 10 minutes at 10,000g). Residual ethanol was removed by air drying ten minutes at 37°C, followed by resuspension of the pellet in 100μL water overnight at 4°C without agitation of any kind. The pooled sample was quantified using the Qubit Broad-Range DNA concentration kit, and was estimated at 323.35 ng/μL with an OD_260/280_ = 1.85. The extracted HMW DNA was used for both short- and long-read sequencing. RNA was extracted from an aliquot of the same fecal sample using PowerMicrobiome RNA isolation kit (cat. no. 26000-50, MoBio) as suggested by the manufacturer. For the protein extractions, a modified protocol based on previously established sequential extraction method [20] was used. Briefly, proteins were precipitated by adding one volume of APP Buffer to the flow-through from an independent RNA purification, followed by mixing and incubation for 10 minutes at room temperature. After incubation, the mixture was centrifuged for 10 minutes at 12000 g and the pellet was washed twice in 70% ethanol, with 1 minute centrifuge cycles at 12000 g, and dried at room temperature for 7 minutes after removing excess ethanol. The pellet was then dissolved in 100μL ALO buffer and incubated for 5 minutes at 95 °C. After complete dissolution and denaturation of the protein, the sample was cooled to room temperature and centrifuged for 1 minute at 12000 g, from which the supernatant was collected for downstream protein analysis.

### Sequencing

#### Short-read sequencing

All DNA samples were subjected to random shotgun sequencing. The sequencing libraries were prepared using Kapa hyperplus Kit (cat. no. 07962401001, Roche) for the fecal sample using the protocol provided with the kit. Enzymatic fragmentation time was 15 minutes to aim for 350bp average size. There was no additional PCR amplification of prepared libraries.

RNA samples for metaT analysis were subjected to rRNA depletion using the QIAseq FastSelect 5S/16S/23S kit (cat. no. 335921, Qiagen) for the fecal sample. Library preparation of rRNA-depleted RNA was done using TruSeq Stranded mRNA library preparation kit (cat. no. 20020594, Illumina) according to the protocol provided by the manufacturer with the exception of omitting the initial steps for mRNA pull down.

Both metaG and metaT libraries were quantified using Qubit HS assay (Invitrogen) and their quality was assessed on a Bioanalyzer HS chip (Agilient). We used the NextSeq500 (Illumina) instrument to perform the sequencing using 2×150bp read length at the LCSB Sequencing Platform.

#### Long-read sequencing

DNA library for the fecal sample was size selected using AMpure beads for longer fragments. The DNA was sheared using a G-tube (cat. no. 520079, Covaris) aiming for 8kb average size according to the protocol provided by the manufacturer. Library preparation for long read sequencing was done using genomic DNA ligation kit (SQK-LSK109) according to the protocol provided. Once all the library loaded on the flowcell was finished, the library was reloaded after either flowcell wash or nuclease flush. In total, the library was loaded 4 times to achieve 16Gbp of sequencing data for this fecal sample.

### Data analysis

Snakemake (v. 5.18.1) [21] was used to implement the analysis workflow. We provide a brief description of the most important steps in the following.

### Sequence data preprocessing

#### Short-reads

The raw short reads were trimmed and preprocessed with fastp (v. 0.20.0) [22] with a minimum length of 40 bp. FastQC (v. 0.11.9) [23] reports were generated from the processed FASTQ files. MetaT short reads from the GDB sample were filtered by discarding reads mapping to rRNA gene references included in the repository of SortMeRNA [24] (v4.2.0-10-g1358b9b, https://github.com/biocore/sortmerna) using BBDuk from the BBMap toolkit (v.38.86, kmer length set to 31 bp) [25]. Additionally, for the GDB sample, reads mapping to the human genome (GCF_000001405.38_GRCh38.p12) were removed using BBDuk (kmer length set to 31 bp, input and output quality encoding offset set to 33).

#### Long reads

For each sample except NWC, single-FAST5 files were converted to multi-FAST5 files using single_to_multi_fast5 from ont-fast5-api (v. 3.1.5), the resulting files were basecalled using guppy on a GPU node (v. 3.6.0+98ff765, configuration file dna_r9.4.1_450bps_modbases_dam-dcm-cpg_hac.cfg, disabled transmission of telemetry pings, chunk size of 1000, 8000 records per FASTQ file) and concatenated into a single FASTQ file. For NWC, no FAST5 were available and, thus, only the provided FASTQ file was used for the analysis. Nanostat (v. 1.1.2) [26] reports were created from the FASTQ files using default parameters. As for the short reads, long reads of the GDB sample were filtered to remove reads mapping to the human genome (GCF_000001405.38_GRCh38.p12) using the same parameters.

### Metagenomic assembly

#### Short-reads

Short-read assemblies were done using preprocessed reads, and MEGAHIT and metaSPAdes. MEGAHIT (v. 1.2.9) [27] was run using default parameters; metaSPAdes (v. 3.14.1) [28] was run using kmer lengths 21, 33, 55 and 77 bp.

#### Long-reads

Long-read assemblies were done using Flye and Raven. Flye (v. 2.8.1) [29] was run by providing the (processed) long reads in a FASTQ file (input parameter “--nano-raw”) and with the flag “--meta”. Raven (v. 1.2.2) [30] was run with default parameters. Assemblies were polished using long and short reads: one round of Racon (v. 1.4.13) [31] with long reads using the flag “--include-unpolished” where reads were mapped to contigs using BWA MEM (v. 0.7.17) [32] with the option “-x ont2d” and processed using samtools (v. 1.9); four rounds of Racon with short reads using the flag “--include-unpolished” where reads were mapped to contigs using BWA MEM and processed using samtools; one round of Medaka (v. 0.8.1) [33] with long reads using the model “r941_min_high”.

#### Hybrid

Hybrid assemblies, i.e. using short and long reads together, were done using metaSPAdes and OPERA-MS. SPAdes was run with the flag “--meta” and the same k-mer lengths as the SR assemblies by additionally providing the long reads using the input parameter flag “-- nanopore”. OPERA-MS (v. v0.8.2-63-gc18b4f3) [15] was run using paired short reads, long reads and the SR assemblies created by MEGAHIT and metaSPAdes, respectively, using minimap2 [34] as the long read mapper. The assemblies were polished by running five rounds of Racon with short reads as described for the LR assemblies. If not stated otherwise, only polished contigs were used for the LR and HY assemblies in the following analysis steps.

### Mapping rate and assembly coverage

For the mapping rate, the used reads were mapped back to the contigs and processed using BWA MEM and samtools in the same fashion as described above when polishing the LR and HY assemblies using Racon. For hybrid assemblies, both long and short reads were mapped to the polished contigs and the BAM files were merged using samtools. For the sample GDB, metatranscriptomic (metaT) short reads were also separately mapped to the (polished) contigs. Mapping statistics were computed from the BAM files using samtools’ options “flagstat”, to determine the number of reads mapping back to the assemblies, and “idxstats” for per-contig mapping information. For GDB, metaT per-base coverage was computed for each assembly from the BAM files using bedtools (v. 2.29.2)[35] (utility “genomecov” with the parameter “-d”).

### Assembly annotation

For each sample and assembly, protein prediction was done using Prodigal (v. 2.6.3) [36] using the option “-p meta”; the keyword “partial” in the headers of the obtained protein FASTA files was used to distinguish complete and partial proteins. Known antibiotic resistance factors were searched in the predicted proteins (after discarding the stop codon symbol “*” from the FASTA files) by running RGI (v. 5.1.1) [37] together with the CARD database (v. 3.1.0) [18] and DIAMOND (v. 0.8.36) [38] for protein alignments. Loose hits flagged as “nudged” by the tool were highlighted as such (i.e. as “nudged”) in the downstream analysis.

The tool barrnap (v. 0.9) [39] was run to predict rRNA genes on assembly contigs using the four provided databases of bacterial, archaeal, metazoan mitochondrial and eukaryotic rRNA genes, respectively. Predictions containing the word “partial” in their product annotation in the obtained GFF files were considered as partial hits.

### Analysis

Assembly statistics were computed by running metaQUAST (v. 5.0.2) [40] without using any genome references, setting the minimum contig length to 0 bps and retrieving the statistics for the contig length thresholds of 0, 1000, 2000 and 5000 bps subsequently. Per sample, assemblies were compared using Mash (v. 2.2.2) [41]: sketches were computed per assembly using a k-mer length of 31 bps and a sketch size of 100000, and pairwise distances were then estimated. Per sample, proteins from all assemblies were clustered using MMseqs2 (v. 12.113e3) [42]. First, a database was created from a concatenated FASTA file of protein sequences (“-- dbtype 1”). Then, option “linclust” with default parameters was used to perform the clustering and the obtained files were converted to tables using option “createtsv“. DIAMOND (v. 0.9.25) [38][43] with the option “blastp” and default parameters was used to align the predicted proteins against the UniProtKB/TrEMBL database (downloaded and created on August 24 2019 from http://ftp.uniprot.org/pub/databases/uniprot/current_release/knowledgebase/complete/, archive uniprot_trembl.fasta.gz) [44]. The created DAA files were converted to tables using option “view” and the parameter “--max-target-seqs 1”. When processing the hits, these were sorted per query and e-value in an ascending order and only the first hit was used. For GDB and metaT, using the per-base coverage information computed for each assembly, average coverage was computed for the corresponding gene sequences of each predicted protein.

### MS/MS acquisition and metaproteomic analysis

1μg of extracted proteins was denatured and briefly loaded on a SDS gel to produce one gel band. The reduction, alkylation and tryptic digestion of the proteins into peptides were performed in-gel. The tryptic peptides were extracted from the gel and desalted prior to mass spectrometry analysis. Peptides were analyzed using a nanoLC-MS/MS system (120 minutes gradient) connected to a Q-Exactive HF orbitrap mass spectrometer (Thermo Scientific, Germany) equipped with a nano-electrospray ion source. The Q-Exactive mass spectrometer was operated in data-dependent mode and the 10 most intense peptide precursor ions were selected for fragmentation and MS/MS acquisition.

For each assembly separately and for all assemblies together, the FASTA file of predicted proteins was concatenated with a cRAP database of contaminants [45] and with the human UniProtKB Reference Proteome prior metaproteomic search. In addition, reversed sequences of all protein entries were concatenated to the databases for the estimation of false discovery rates. The search was performed using SearchGUI-3.3.20 [46] with the X!Tandem [47], MS-GF+ [48] and Comet [49] search engines and the following parameters: Trypsin was used as the digestion enzyme, and a maximum of two missed cleavages was allowed. The tolerance levels for matching to the database was 10 ppm for MS1 and 0.2 Da for MS2. Carbamidomethylation of cysteine residues was set as a fixed modification and protein N-terminal acetylation and oxidation of methionines was allowed as variable modification. Peptides with length between 7 and 60 amino acids, and with a charge state composed between +2 and +4 were considered for identification. The results from SearchGUI were merged using PeptideShaker-1.16.45 [50] and all identifications were filtered in order to achieve a protein false discovery rate (FDR) of 1%.

### Plots

Figures were generated in R (v. 4.0.2, https://www.r-project.org/) using, *inter alia*, Pheatmap (v. 1.0.12, https://github.com/raivokolde/pheatmap) for heatmap plots, UpSetR (v. 1.4.0) [51] for intersection plots, ggplot2 (v 3.3.2) [52] and its various extensions for other plot types, color palettes from the viridis (v. 0.5.1, developed by Stéfan van der Walt and Nathaniel Smith, https://github.com/sjmgarnier/viridis) and ggsci (v. 2.9, https://github.com/road2stat/ggsci) packages and the patchwork package (v. 1.1.1, https://github.com/thomasp85/patchwork) for combining plots.

---

**Supp. Fig. 1:**
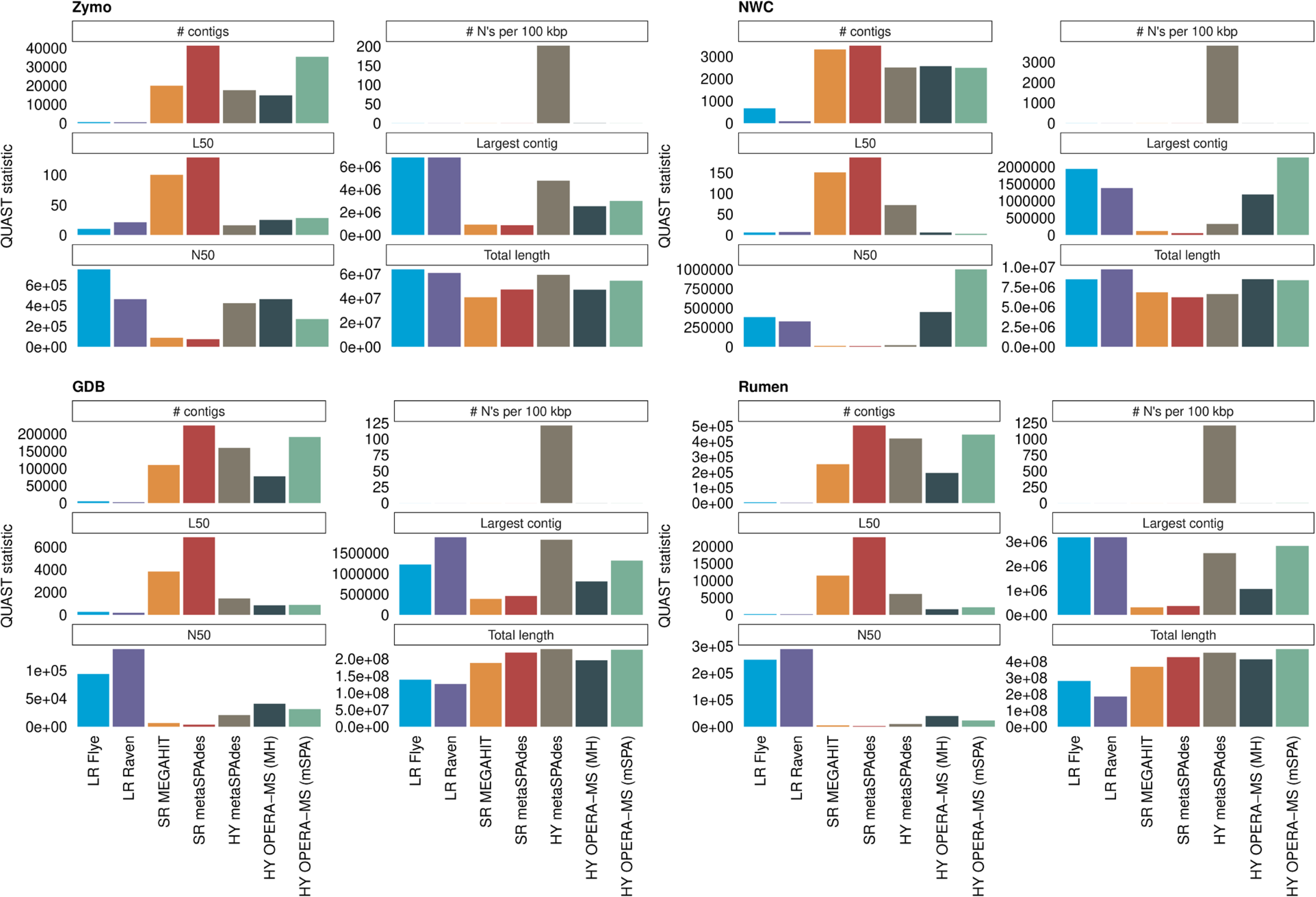
Assembly statistics. Assembly statistics for each assembly and sample including the total number of contigs, number of N bases per 100kbp, L50 value (number of contigs), N50 value (in bps), the length of the largest contig (in bps), and the total assembly length (in bps). The color corresponds to the tool used for metagenomic assembly.

**Supp. Fig. 2:**
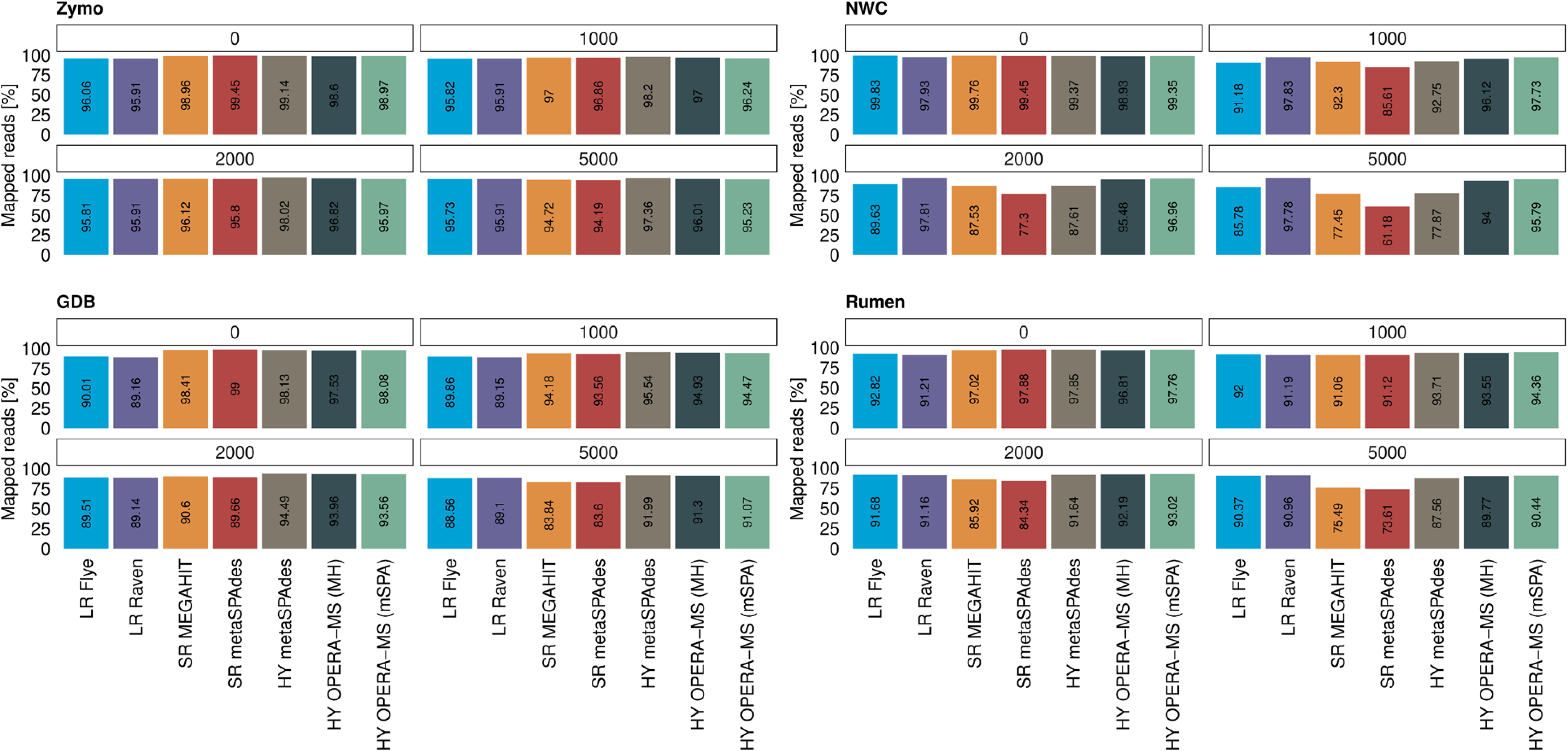
Mapping rate of metagenomic reads. Mapping rate of metagenomic reads to each assembly for each sample considering all contigs and contigs being at least 1000, 2000 and 5000bps long. The color corresponds to the tool used for metagenomic assembly.

**Supp. Fig. 3:**
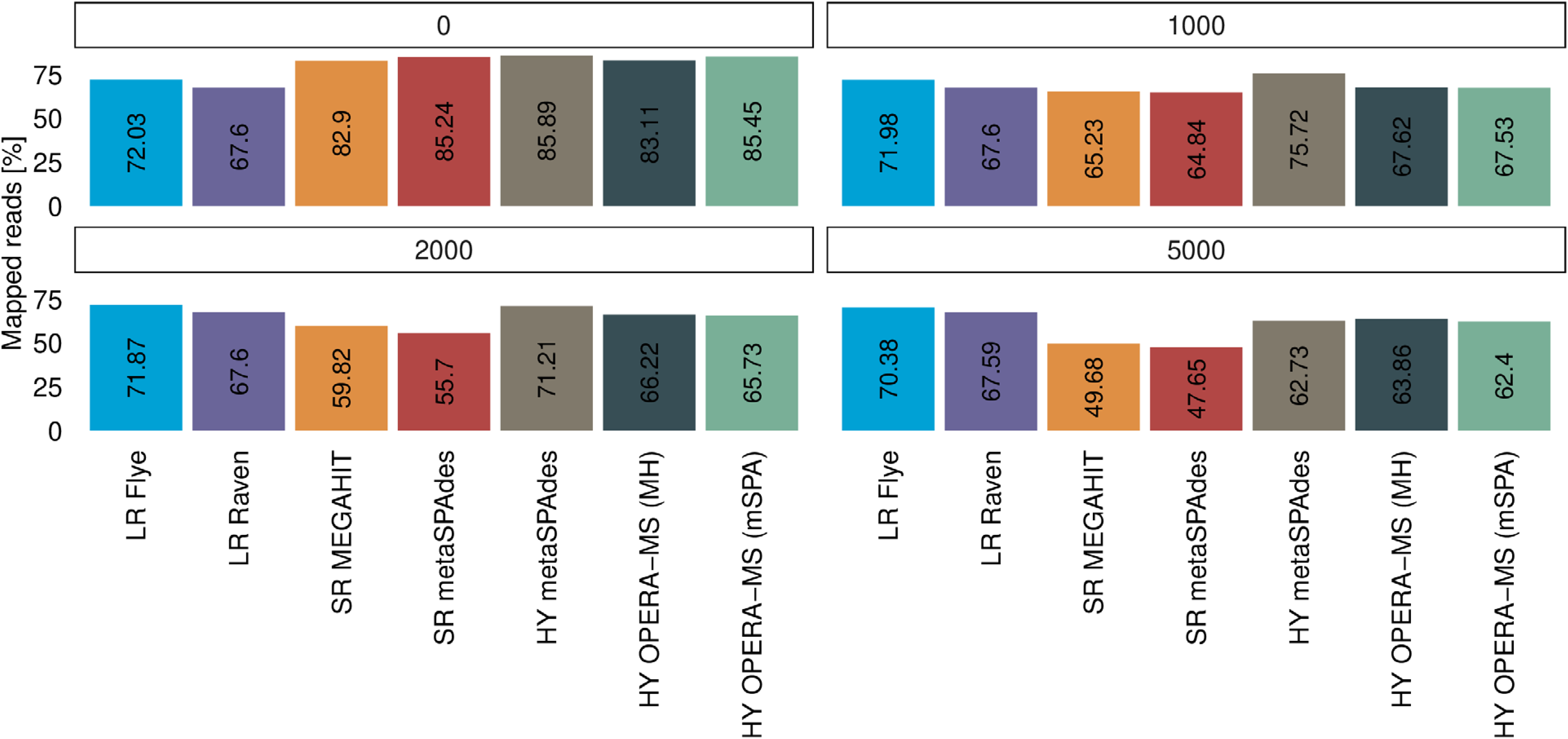
Mapping rate of metatranscriptomic reads. Mapping rate of metatranscriptomic reads to each assembly in GDB considering all contigs and contigs being at least 1000, 2000 and 5000bps long. The color corresponds to the tool used for metagenomic assembly.

**Supp. Fig. 4:**
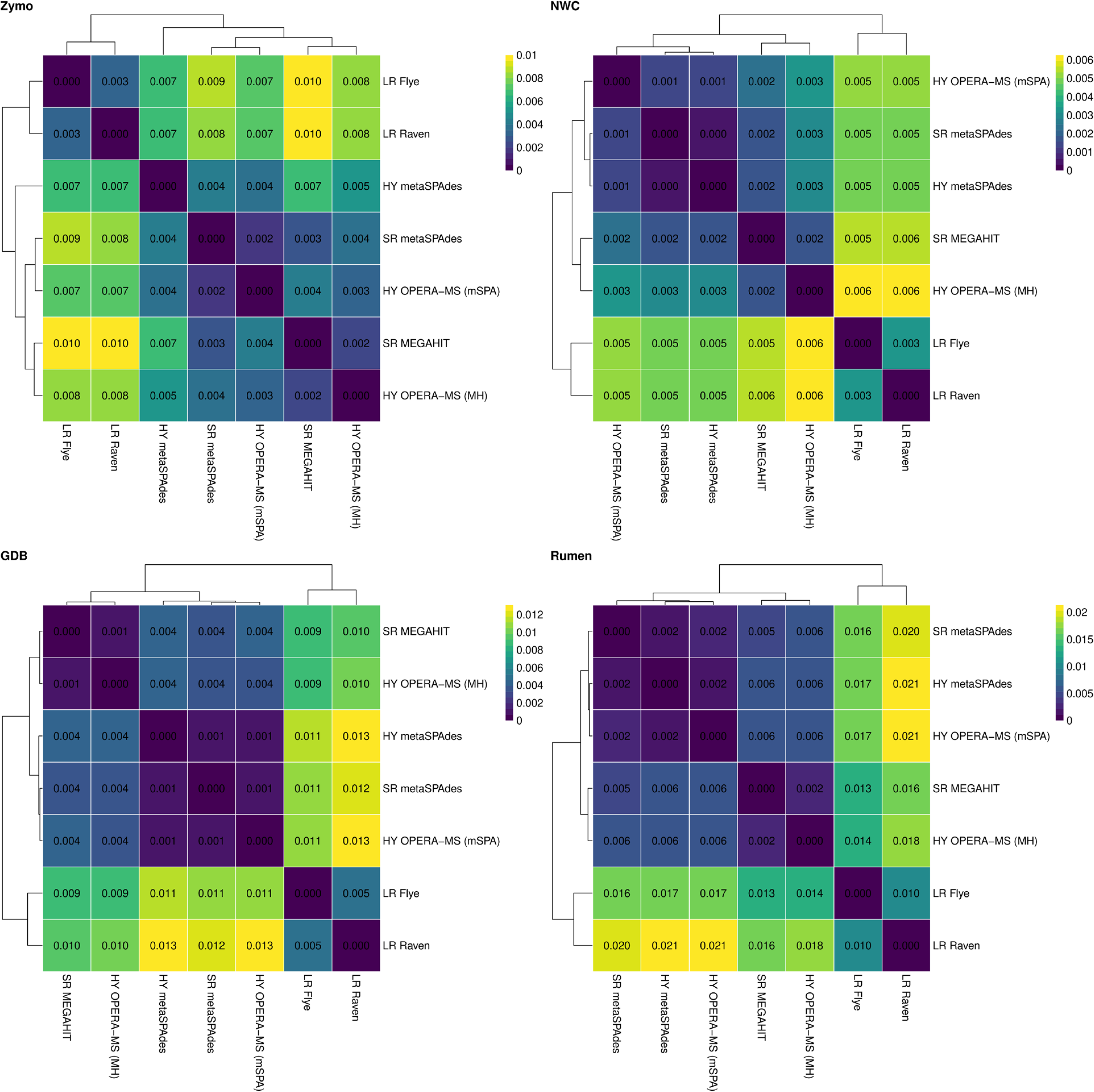
Assembly similarity. Heatmap of assembly dissimilarity of each sample. The cell color corresponds to the estimated dissimilarity value and the rounded value is shown in each cell: higher values (yellow) indicate higher dissimilarity, lower values (dark purple) indicate high similarity. Assemblies were grouped using hierarchical clustering (linkage method “complete”): the resulting trees are shown in the heatmaps.

**Supp. Fig. 5:**
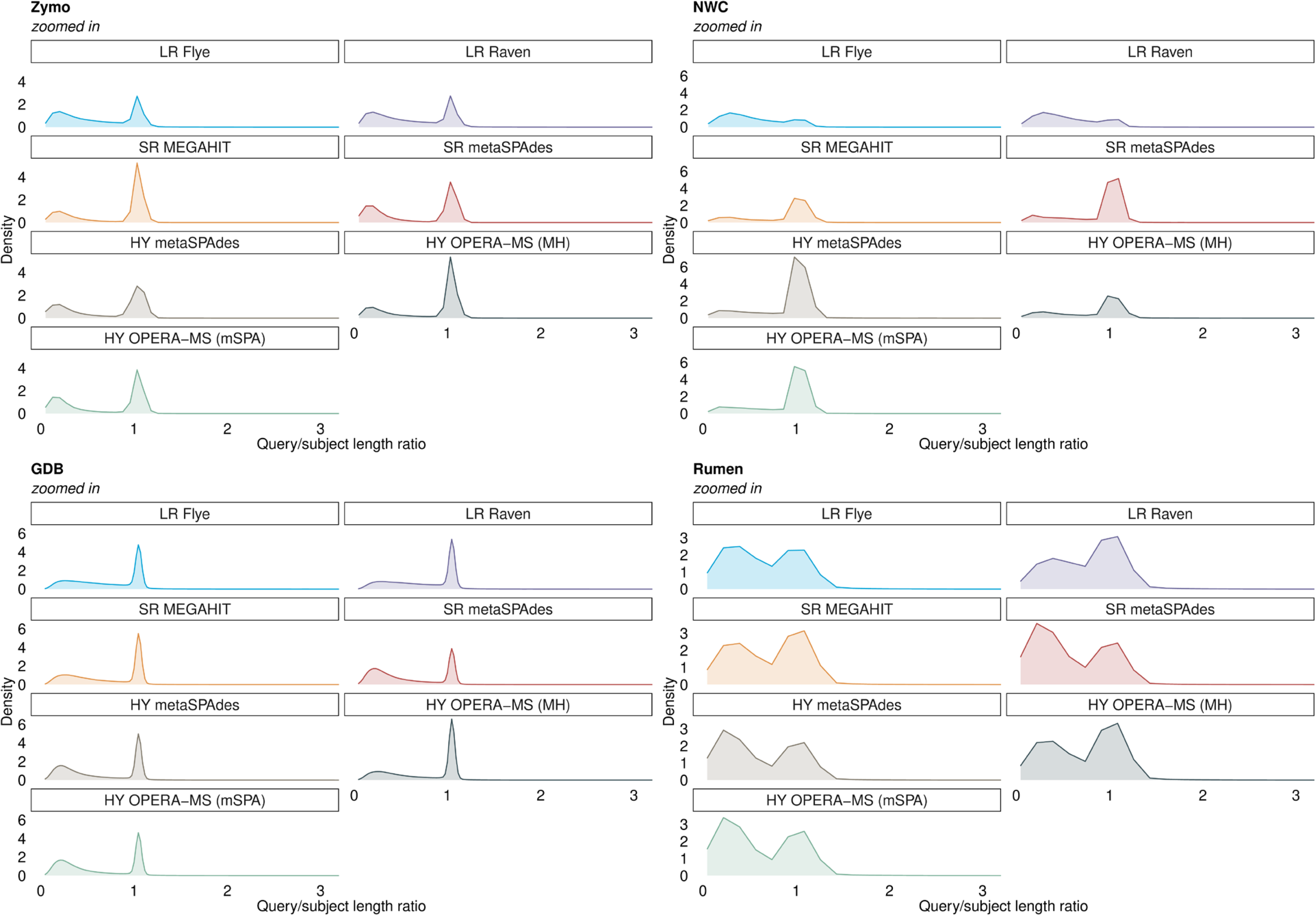
Protein sequence search in the UniProtKB/TrEMBL nr database. Density distribution of the query/subject length ratios of the best hit obtained in the protein sequence search in the UniProtKB/TrEMBL nr database for each assembly and sample. The color corresponds to the tool used for metagenomic assembly.

**Supp. Fig. 6:**
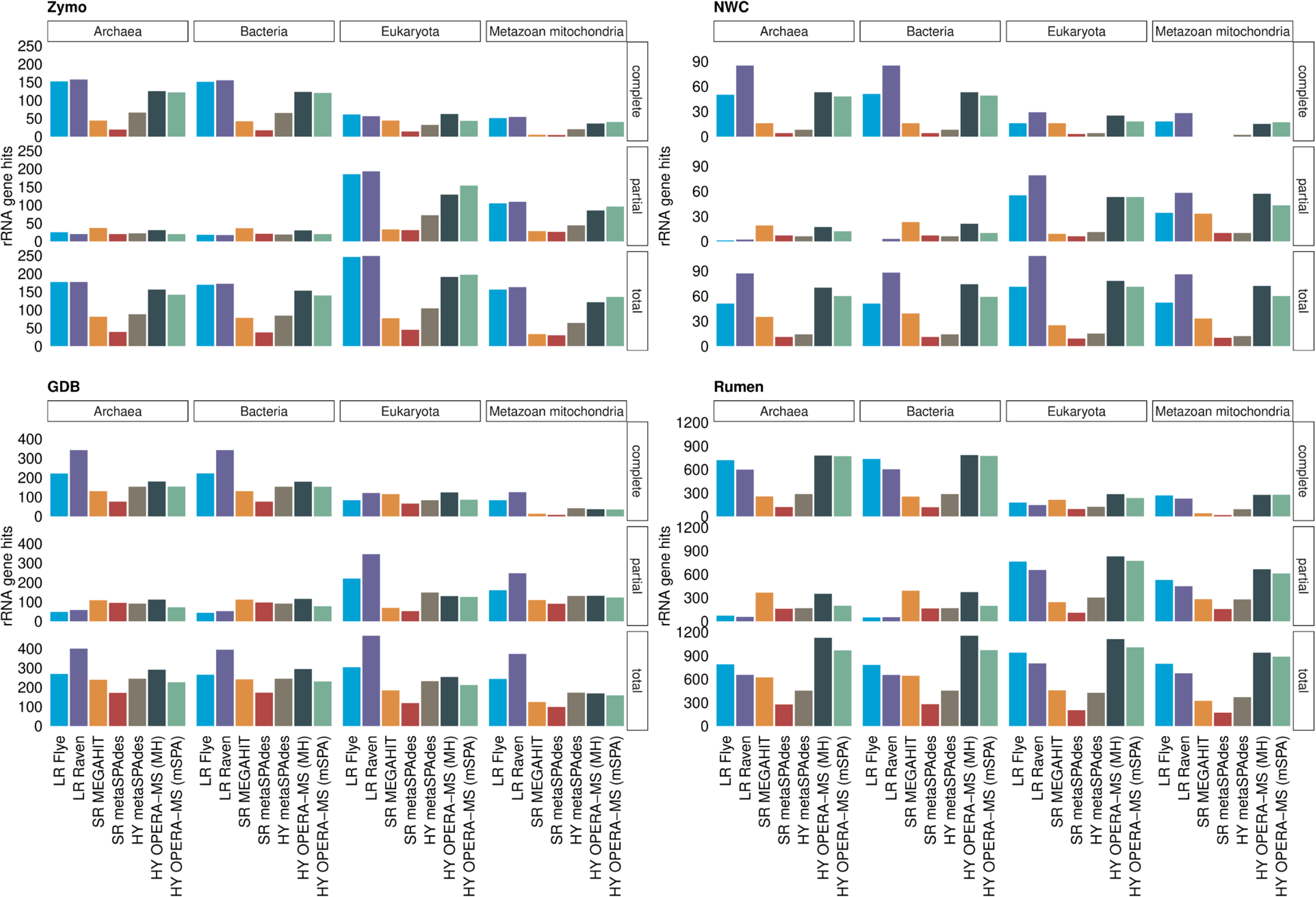
Prediction of rRNA genes. Number of rRNA genes (complete, partial and total) found by barrnap in assembly contigs using different rRNA gene databases (Archaea, Bacteria, Eukaryota and Metazoan mitochondria) for each assembly and sample. The color corresponds to the tool used for metagenomic assembly.

**Supp. Tab. 1:**
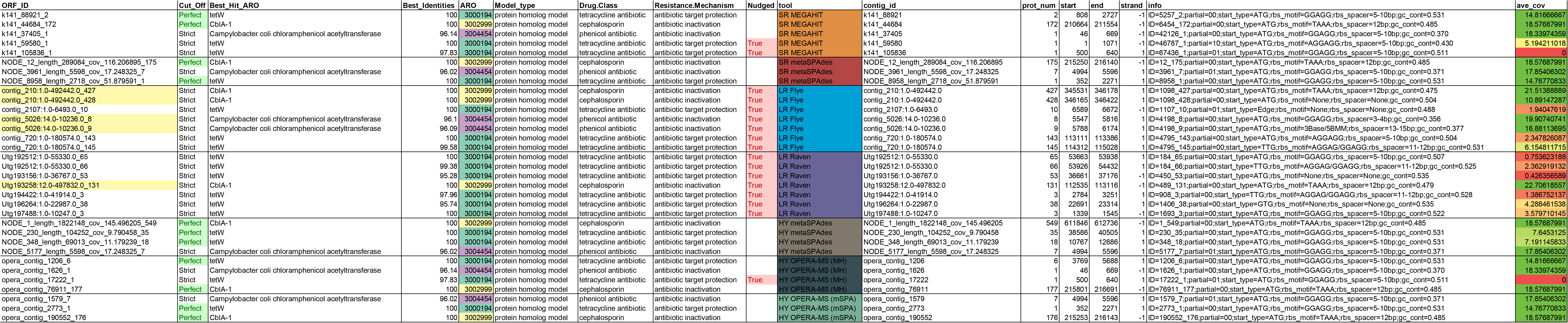
Proteins assigned to exclusive AROs in GDB. Information on proteins from GDB assemblies assigned to AROs found exclusively in SR and HY assemblies when considering only “strict” RGI hits, i.e., AROs 3000194, 3002999 and 3004454. The table includes the protein ID, the RGI hit information (RGI Detection Paradigm, ARO term of top hit in CARD, percent identity of match to top hit in CARD, ARO ID, CARD detection model type, ARO’s drug class and mechanism, flag whether the hit was “nudged” from “loose” to “strict”), the assembly tool, additional protein information (contig ID, protein number on the source contig, start and end coordinates on the contig, Prodigal’s annotation information) and average metaT coverage.

